# PopTradeOff: a database for exploring population-specific trade-offs between adaptive evolution, disease susceptibility, and drug responsiveness

**DOI:** 10.1101/2023.02.09.527958

**Authors:** Ji Tang, Huanlin Zhang, Hai Zhang, Hao Zhu

## Abstract

The influence of adaptive evolution on disease susceptibility has drawn attention, but the extent of the influence, whether favored mutations also influence drug responses, and whether the associations between the three are population specific remain little known. Using a deep learning network to integrate seven statistical tests for detecting selection signals, we predicted favored mutations in the genomes of 17 human populations. We integrate these favored mutations with GWAS sites and drug response-related variants into the database PopTradeOff. The database also contains genome annotation information on the SNP, sequence, gene, and pathway levels. The preliminary data analyses suggest that substantial associations exist between adaptive evolution, disease susceptibility, and drug responses. The database may be valuable for disease studies, drug development, and personalized medicine.

## 1. Introduction

Abundant data generated by genome-wide association studies (GWAS) and clinical drug research have revealed the population-specificity of many GWAS sites and drug response-related sites (Beck et al., 2020; Buniello et al., 2019; Whirl-Carrillo et al., 2021; Wishart et al., 2006). How population-specificity is determined, and especially whether disease susceptibility and drug responsiveness have intrinsic associations, is important for basic, translational, and clinical biomedical research. A plausible supposition is that both are related to the evolution of specific adaptations in human populations. The supposition is supported by findings such as genetic ancestry plays a central role in population pharmacogenomics (Yang et al., 2021). However, few systematic investigations have been reported.

During and after migration out of Africa, many single nucleotide mutations occurred in the genomes of different populations, adapting these populations to different environments, climates, and lifestyles (Figure 1A) (1000 Genomes Project et al., 2015) (Fan et al., 2016; Jeong and Di Rienzo, 2014). Thus, these mutations are beneficial (favored) and influence phenotypic differences between human populations. However, recently, evidence has emerged suggesting that many favored mutations have been selected with a cost; that is, they also make the carriers susceptible to certain diseases (Benton et al., 2021; Prohaska et al., 2019; Tang et al., 2022). Increasing findings indicate that favored mutations are both population- and disease-associated. For example, the mutations in the *HBB* gene make some Africans resistant to P. falciparum malaria but also susceptible to sickle cell anemia (Ackerman et al., 2005), and the mutations in genes encoding hypoxia-inducible factors (HIFs) make the carriers benefit from increased oxygen delivery but also suffer from increased blood viscosity (a contributing factor to the high incidence of stroke in Tibetans (GBD 2019 Stroke Collaborators, 2021). These findings provide the primary support for the supposition that population-specific favored mutations critically associate the triangle relationships between adaptive evolution, disease susceptibility, and drug responsiveness (Figure 1B). Deciphering the association helps us understand human evolution and improves personalized medicine.

**Figure 1.**
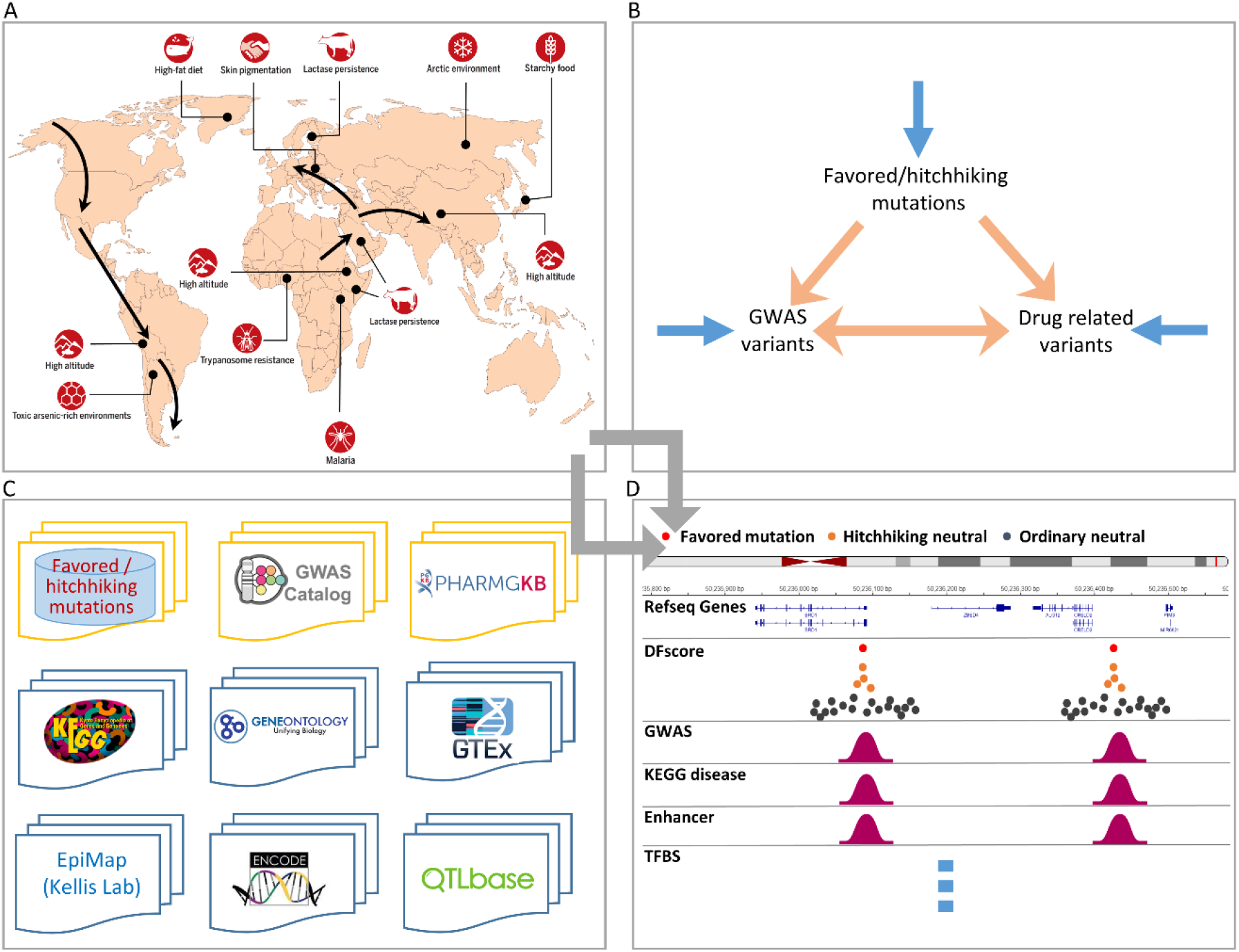
An overview of PopTradeOff. (A) During and after the out-of-Africa migration, many population-specific mutations occurred and made humans adapt to varied local environments and changed lifestyles but also susceptible to particular diseases (adapted from (Benton et al., 2021; Fan et al., 2016)). (B) PopTradeOff helps examine the associations between the three families of mutations. Blue arrows indicate the traditional studies, including population genetics, GWAS study, and pharmacogenetics. Orange arrows indicate the associated studies. (C) PopTradeOff contains data from multiple sources. (D) An illustration of the genome browser page that displays a search result in tracks.

Identifying favored mutations has been a challenge for decades. Many statistical tests have been developed to detect signals generated by favored mutations (especially selective sweeps) (Vitti et al., 2013). However, detecting favored mutations per se is much more difficult, and integrating multiple statistical tests is a highly recognized strategy (Grossman et al., 2013; Sabeti et al., 2006). Recently, we used a deep learning network (called DeepFavored) to integrate seven statistical tests to systematically identify favored mutations in the human population CEU, CHB, and YRI (Tang et al., 2022). DeepFavored outperforms other integration methods, and the combined analysis of favored mutations and GWAS sites reveals the intrinsic association (i.e., extensive trade-offs) between population-specific adaptive evolution and disease susceptibility (Tang et al., 2022).

In this study, we further trained the DeepFavored network using the demographic models of 17 populations and used the trained network to identify favored mutations in the 17 populations. Further, we integrated favored and hitchhiking neutral mutations (hereafter just called hitchhiking mutations) with disease/trait phenotype and drug response phenotype data into the database PopTradeOff (Figure 1C). This database helps unveil the associations between adaptive evolution, disease susceptibility, and drug responsiveness for studying human physiology and diseases and performing precision and personalized medicine. Preliminary data analyses indicate that many favored and hitchhiking mutations influence gene expression and function, disease susceptibility, and drug responses population-specifically (Figure 1D).

## 2. Results

### 2.1. Database overview

The trained deep learning network DeepFavored outputs a score (called DFscore) for each mutation in the genome of each population and a rank of the score in a 600 KB window centered on the mutation (Figure 1) (Tang et al., 2022). DFscore >= 0.5 and rank <= 3 strongly suggest favored mutations. Upon the stringent criteria DFscore >= 0.5 and rank <= 3, 21293 favored mutations were identified in the 17 populations, and the probability a neutral mutation meets the criteria is <1e-6.

Then, in windows containing favored mutations, 867379 neutral mutations having LD > 0.6 with favored mutations were identified as hitchhiking mutations (Figure 2A). Many favored and hitchhiking mutations overlap GWAS sites in GWAS Catalog and drug response-related sites in DrugBank and PharmGKB (Figure 2B). The functional significance of population-specific favored and hitchhiking mutations was examined upon multiple data families (e.g., QTLbase, EpiMap, KEGG), and many mutations were identified in regulatory and functional sites (Figure 2C). Of note, 5353 (25.14%) favored mutations and 178568 (20.59%) hitchhiking mutations overlap GWAS sites, and many overlapping sites are associated with skin, nervous, immune, and metabolic system diseases (Figure 2D). It is found that the influence of favored and hitchhiking mutations on gene expression is highly tissue-specific (Figure 2E). These results indicate that adaptive evolution considerably influences gene expression and disease susceptibility.

**Figure 2.**
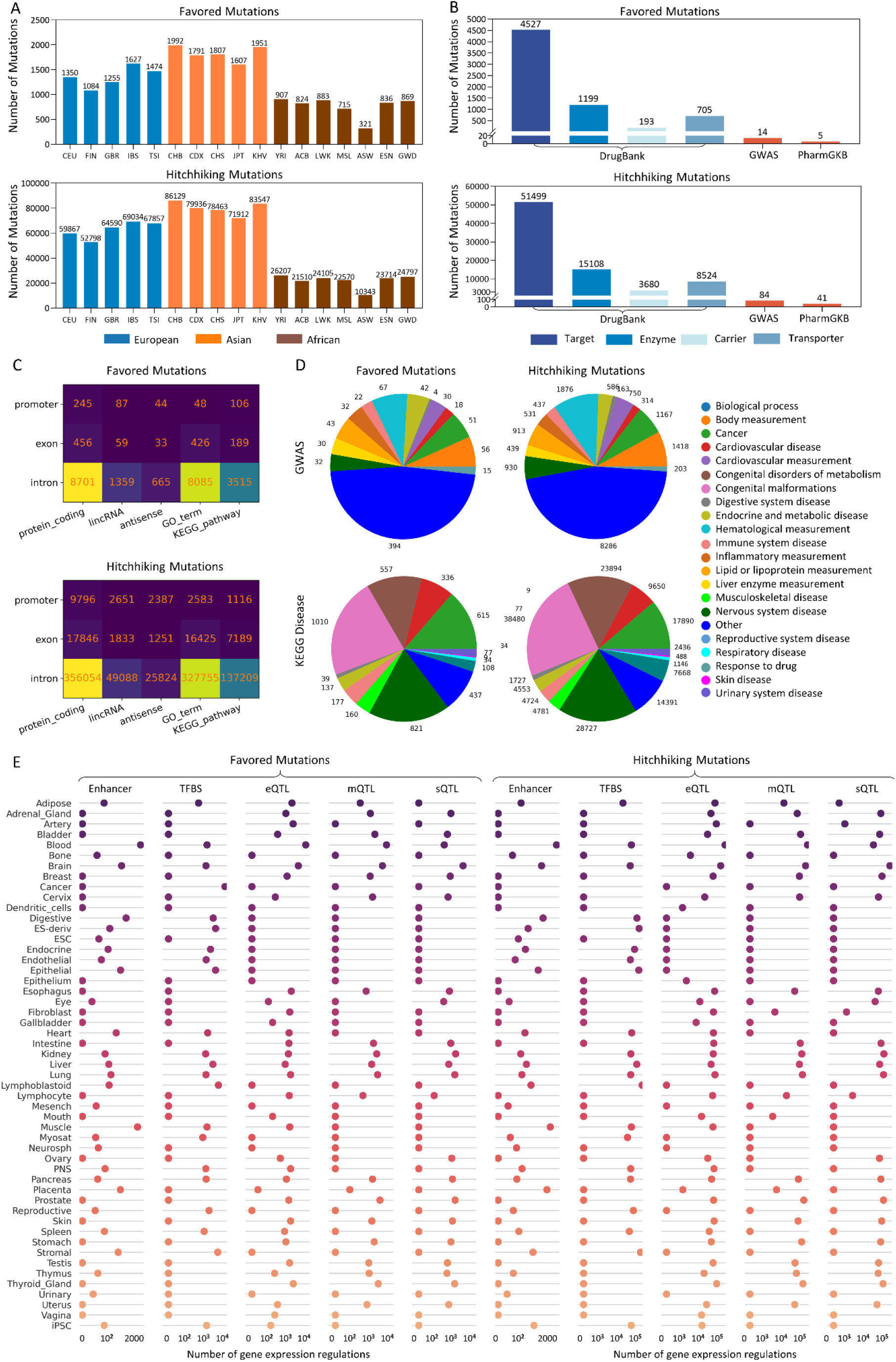
Data summary of PopTradeOff. (A) The number of favored and hitchhiking mutations in 17 European, Asian, and African populations. (B) The number of favored and hitchhiking mutations that overlap GWAS and drug response-related sites. Drug response-related genes in DrugBank were categorized into four classes: Target, Enzyme, Carrier, and Transporter. (C) The number of favored and hitchhiking mutations located in different genomic regions, gene types, gene sets, and pathways. (D) The number of favored and hitchhiking mutations potentially related to different diseases. (E) Favored and hitchhiking mutations at different regulatory loci influence tissue-specific gene expression. The horizontal axes denote the number of genes at these regulatory loci and in specific tissues.

**Figure 3.**
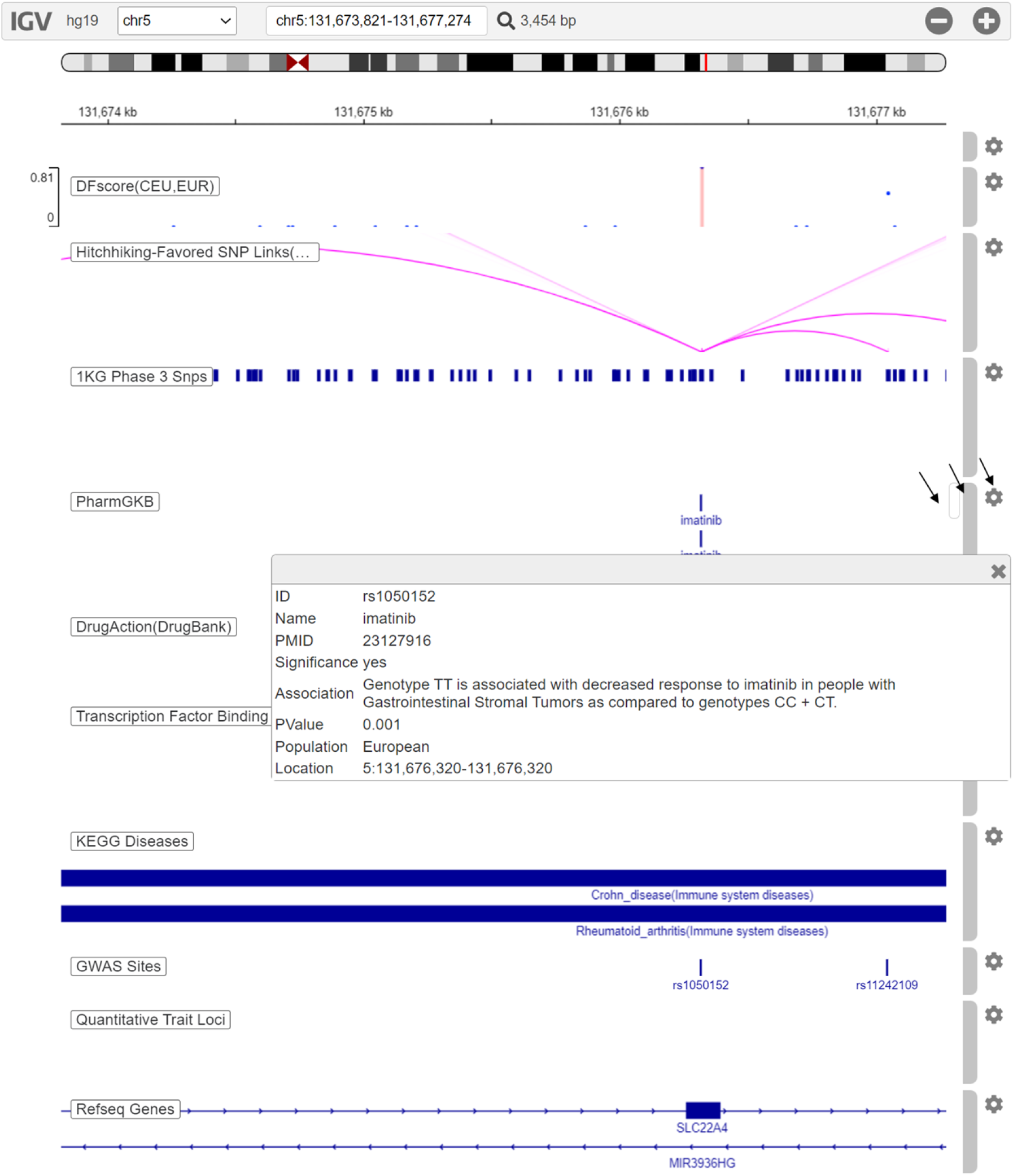
The genome browser webpage shows some tracks of rs1050152. Here rs1050152 is a search result under the combined conditions (1) “Specify a mutation type” =“ Favored”, (2) “Specify a population” =“ All”, (3) “Input a KEGG disease ID or a keyword” =“ crohn”. The left side displays track names. On the right side are track-specific vertical scroll bars (each track has its own scroll bar for displaying multiple sub-tracks), track highlighting bars, and track configuration buttons (indicated by arrows). The “DFscore” track shows that the mutation has a high DFscore in the population CEU and super-population EUR. The “Hitchhiking-Favored SNP Links” track shows the hitchhiking mutations of this mutation. The “1KG Phase 3 Snp” track shows the SNPs of 1000 Human Genome. Clicking an “imatinib” record in the PharmGKB track, a window pops up and displays the content of the PharmGKB record. The “KEGG Diseases” track shows that this mutation is involved in Crohn’ s disease and Rheumatoid arthritis. The “GWAS Sites” track indicates this mutation is a GWAS site. This example demonstrates the associations between adaptive evolution, disease susceptibility, and drug responsiveness.

Favored and hitchhiking mutations that overlap drug response-related sites are limited. However, many favored and hitchhiking mutations are found in exons or QTLs of drug response-related genes, suggesting that adaptive evolution may influence drug responses on the gene level.

### 2.2. Key functions

#### Search engine

A search engine is developed for searching favored mutations, hitchhiking mutations, GWAS sites, and drug response-related sites. By searching data upon flexibly defined conditions, the database can be used to identify (i) favored and hitchhiking mutations and genes influenced by adaptive evolution in the 17 human populations, (ii) whether these genes influence drug responses and actions in a population-specific manner, (iii) whether these genes influence the susceptibility/severity of specific diseases, (iv) the distribution of disease- and drug-related favored/hitchhiking mutations in specific genomic regions. The results of a search are displayed in a Result table.

#### Genome browser

To look into details of a record in the result table, click that record to open the genome browser. The graphical genome browser displays a mutation and relevant attributes in tracks in the context of the human genome hg19. The genome background shows the information that a mutation plays its role. The order, height, and color of tracks can be configured flexibly. Clicking the stripes, points, or arcs in the tracks can open popup windows to show more details.

#### Drug response information

Some mutations have many drug response-related records in PharmGKB and DrugBank. To display details of these record, clicking the record names in the PharmGKB and DrugBank tracks to open popup windows.

### 2.3. Cases of associations between adaptive evolution, disease susceptibility, and drug responses

Here we present several examples to illustrate using PopTradeOff to explore the associations between adaptive evolution, disease susceptibility, and drug responses.

#### The immune disease-related rs1050152 hosts a favored mutation in European populations

rs1050152 is an immune disease-related variant in some European populations. The related diseases include Crohn’s disease, asthma, and ulcerative colitis (https://www.gwascentral.org/marker/HGVM236012/results). It is also a drug response-related variant, as genotype CT is associated with increased response to Ustekinumab in people with Psoriasis as compared to genotype CC (p-value = 0.037), and genotype TT is associated with decreased response to Imatinib in people (especially Europeans) with gastrointestinal stromal tumors, leukemia, and myelogenous as compared to genotypes CC + CT (https://www.pharmgkb.org/variant/PA166156750/variantAnnotation). The deep learning network identifies the mutation T on rs1050152 as a strongly favored mutation in CEU (DFscore = 0.815). Ustekinumab is used to treat immune-related diseases (e.g., plaque psoriasis and Crohn’s disease), and Imatinib is used to treat certain types of leukemia, skin cancer, and gastrointestinal stromal tumors. The predicted favored mutation T, together with the GWAS and PharmGKB data, highlight that this mutation may have been selected for with changed susceptibility of these immune-related diseases and altered responsiveness to the related drugs.

#### The scalp hair shape-related rs11150606 hosts a favored mutation in East Asians

rs11150606 is a phenotype-related variant in East Asian populations. The related phenotype is scalp hair shape (https://www.gwascentral.org/marker/HGVM3858600/results). It is also a drug response-related variant, as allele C is associated with a decreased dose of Warfarin as compared to allele T (p = 0.0221) (https://www.pharmgkb.org/variant/PA166155104/variantAnnotation). Warfarin is used to prevent blood clots from forming or growing larger in blood and blood vessels. The mutation C on rs11150606 has a high frequency (>76.4%) in East Asian populations but a low frequency (< 20.1%) in other populations. The deep learning network identifies the mutation C on rs11150606 as a strongly favored mutation in East Asians (DFscore = 0.713/0.795/0.651/0.667 in CDX/CHS/JPT/KHV, respectively). The predicted favorable mutations together with the GWAS and PharmGKB data highlight that positive selection for the favorable mutation C may be associated with scalp hair shape and warfarin responsiveness.

#### rs776746 contributes greatly to inter-racial variation in drug metabolism

rs776746 is an SNP encoding the (nonfunctional) CYP3A5*3 allele of the *CYP3A5* gene. Because the *CYP3A5* gene encodes a member of the cytochrome P450 superfamily of enzymes, the cytochrome P450 proteins catalyze the metabolism of many drugs and the synthesis of cholesterol, steroids, and other lipids, CYP3A5 is an important contributor to inter-racial variation in drug metabolism. Of note, the frequency of the C allele is low in AFR but high in all other populations (especially in Europeans), and the frequency of the T allele shows the opposite (high in AFR but low in all other populations, especially in Europeans). While the related GWAS phenotypes are limited, including “glycated hemoglobin levels” and “Tacrolimus trough concentration in kidney transplant patients” (https://www.gwascentral.org/marker/HGVM696974/results), many studies reported rs776746 as a drug response-related variant (https://www.pharmgkb.org/variant/PA166157267/variantAnnotation). Here are some important cases. (i) Genotype CC is associated with decreased dose (or increased risk) of Tacrolimus in people with kidney transplantation as compared to genotypes CT + TT (p < 0.001), and genotype TT is associated with an increased dose of Tacrolimus in people with kidney transplantation as compared to genotype CC (p < 0.0001) (Niioka et al., 2012; Satoh et al., 2009; Vannaprasaht et al., 2013). (ii) Genotype CC is associated with increased metabolism of amlodipine in healthy individuals compared to genotypes CT + TT (Kim et al., 2006). (iii) Genotypes CC + CT are associated with increased response to Atorvastatin compared to genotype TT (Kivisto et al., 2004; Ramakumari et al., 2018; Thompson et al., 2005). (iv) Genotype CC is associated with a decreased dose of cyclosporine compared to genotypes CT + TT (Zhu et al., 2011). (v) Genotype TT is associated with an increased risk of kidney diseases when treated with Dihydropyridine derivatives compared to genotypes CC + CT (Turkmen et al., 2022). The deep learning network identifies the mutation T on rs776746 as a favored mutation in the European population IBS.

## 3. Materials and Methods

### 3.1. Identifying favored mutations in 17 populations

First, we downloaded the genome-wide mutation data (excluding the Y chromosome) of 17 European, Asian, and African populations from 1000 Genomes Project (Phase 3) (1000 Genomes Project et al., 2015). The European populations include Utah residents with Northern and Western European ancestry (CEU), Finnish in Finland (FIN), British in England and Scotland (GBR), Iberian populations in Spain (IBS), and Toscani in Italia (TSI). The Asian populations include Han Chinese in Beijing (CHB), Chinese Dai in Xishuangbanna (CDX), Southern Han Chinese (CHS), Japanese in Tokyo (JPT), and Kinh Vietnamese (KHV). The African populations include Yoruba in Ibadan, Nigeria (YRI), African Caribbean in Barbados (ACB), Luhya in Webuye, Kenya (LWK), Mende in Sierra Leone (MSL), African Ancestry in Southwest US (ASW), Esan in Nigeria (ESN), Gambian in Western Division, and The Gambia (GWD).

Second, we used the demographic models and population genetics data of the 17 populations to train the deep learning network DeepFavored (Tang et al., 2022). The ID lists of unrelated individuals were acquired from the Ensembl website (Cunningham et al., 2022). The trained network integrates seven statistical tests (Fst, XPEHH, iHS, ΔDAF, nSL, iSAFE, and ΔiHH) to identify favored and hitchhiking mutations in the 17 populations (Tang et al., 2022).

Third, favored and hitchhiking mutations in each population were predicted upon scores that reflect the strength of positive selection (called DFscore), together with the ranks of the scores in the 600 KB genomic regions. Previous studies suggest that 600 KB is the size of the genomic region affected by a moderately strong selection (Sabeti et al., 2006). Mutations with DFscore >= 0.5 and rank <= 3 (in each 600 KB region) were identified as favored mutations, and the nearby neutral mutations (within the 600 KB region) with the linkage disequilibrium (LD) > 0.6 with the favored mutation were identified as hitchhiking mutations. Analysis using simulated data indicates that the probability that a neutral mutation gets a DFscore > 0.5 is <1e-6. Finally, correlation coefficients were calculated to measure the likelihood that favored alleles and the alleles in hitchhiking sites are on the same haplotypes.

### 3.2. Integrating favored/hitchhiking mutations with trait/disease and drug response data

We downloaded favored/hitchhiking mutation-related disease/trait and drug response data from GWAS Catalog (ebi.ac.uk/gwas/) (Buniello et al., 2019), PharmGKB (https://www.pharmgkb.org/) (Whirl-Carrillo et al., 2021; Whirl-Carrillo et al., 2012), and DrugBank (https://go.drugbank.com/) (Wishart et al., 2006). GWAS Central also contains many GWAS sites and can be used as an external data source (https://www.gwascentral.org/) (Beck et al., 2020).

### 3.3. Collecting genome annotation data

Genome annotation data include the Ensembl genome annotation (GRCh37 release 87) downloaded from the Ensembl website (Cunningham et al., 2022), the pathway information downloaded from the KEGG website (Kanehisa and Goto, 2000), and the gene set information downloaded from the Gene Ontology (GO) website (Ashburner et al., 2000; Gene Ontology, 2021). Disease annotation data were downloaded from the KEGG Disease database (Kanehisa and Goto, 2000). Quantitative trait loci data across multiple human molecular phenotypes were downloaded from the QTLbase website (Zheng et al., 2020). The annotations of enhancers, chromatin states, upstream regulators, and downstream target genes upon compiling 10,000 epigenomic maps across 800 human samples were downloaded from the EpiMap website (Boix et al., 2021). The transcription factor (TF) binding sites (TFBS) data, which contain 338 TFs and 130 cell types generated by the ENCODE project (Encode Project Consortium, 2012), were downloaded from the UCSC Genome Browser (http://genome.ucsc.edu/). For each TFBS, the nearest genes were treated as the target genes of the TF, and the 130 cell types were classified into 33 groups upon the *main_metadata_table*.*tsv* in the EpiMap website. The four families of data (favored/hitchhiking mutations, GWAS sites, drug response-related sites, genome annotation data) were integrated into PopTradeOff database using the MySQL software.

### 3.4. Developing the data search and display functions

The user interface of PopTradeOff was built using the Django (https://docs.djangoproject.com/en/4.1/) and VUE frameworks (https://vuejs.org/guide/introduction.html). Data in PopTradeOff can be searched for upon flexibly combined conditions. The igv.js software was revised and used to develop the genome browser that graphically displays search results in UCSC Genome Browser-like tracts (Figure 1D) (Robinson et al., 2020).

## 4. Discussion

Multiple databases, including dbPHSP (Li et al., 2014), 1000 Genomes Selection Browser 1.0 (Pybus et al., 2014), and PopHumanScan (Murga-Moreno et al., 2019), have been developed for analyzing selection signals in human populations. However, the favored mutations that cause these selection signals matter more for precision and personalized medicine, and most favored mutations in human populations remain unknown. Here we report the development of such a database that contains predicted favored mutations in 17 human populations and the integration of the database with GWAS data, drug response-related data, and genome annotation data. A trade-off between adaptive evolution and disease susceptibility has drawn researchers’ attention (Benton et al., 2021; Prohaska et al., 2019). Our previous analyses of favored and hitchhiking mutations and GWAS sites suggest that the trade-off is extensive and population-specific (Tang et al., 2022). Disease-related favored mutations especially influence metabolism, immune, and nervous system diseases, and these systems have evolved substantially during human evolution. The preliminary analyses of PopTradeOff suggest that the association extends to population-specific drug responsiveness. All of these findings point to the necessity of identifying favored mutations in more populations and integrating data from multiple sources. Exploring and understanding the associations between the three aspects may greatly promote disease studies, drug development, and personalized medicine.

Although we have identified substantial overlap between favored/hitchhiking mutations and GWAS sites, the overlaps between favored/hitchhiking mutations and drug response-related variants reported in PharmGKB are limited. Probably more drug response-related variants are to be identified. On the other hand, many favored/hitchhiking mutations are found in exons or QTLs of drug response-related genes, suggesting associations at the gene level.

The favored mutations in the 17 populations may likely be underestimated because we were prudential to adopt the stringent criteria for detecting favored and hitchhiking mutations. Future versions of PopTradeOff will report more favored and hitchhiking mutations, together with updated data from the related sources.

## Acknowledgments

This work was supported by the National Natural Science Foundation of China (31771456).

## Author contribution

J.T. and Hao Zhu designed the study. J.T., Huanlin Zhang, and Hai Zhang built the database and the website. J.T and Hao Zhu wrote the manuscript.

